# *Escargot* is involved in labial and antennal imaginal disc development through two different developmental pathways

**DOI:** 10.1101/2020.02.10.942862

**Authors:** Fernando Rosales-Bravo, Iván Sánchez-Díaz, Enrique Reynaud, Verónica Narváez-Padilla

## Abstract

In insects, imaginal discs form the adult structures. Imaginal discs are formed by two epithelial layers, the lower disc proper columnar epithelium and the upper peripodial squamous epithelium (also known as peripodial membrane). During morphogenesis and metamorphosis there is a complex crosstalk between these two epithelia that defines the final size and form of the adult organs. In this work we found that in the antennal disc, the dosage of the transcriptional factor Escargot (Esg) regulates the extension of the peripodial epithelium. A reduction in Esg expands the peripodial domain at the expense of the antennal disc proper causing a distortion of the anteroposterior compartments resulting in malformations or duplications of antennae and maxillary palps. In the labial disc, a different morphogenetic pathway controls its development, and loss of *esg* produces a complete loss of the proboscis through a pathway that involves *dpp*.

**Summary statement:** The gene *escargot* regulates proboscis, maxillary palps and antennae development in *Drosophila melanogaster* through two different developmental pathways: one involving cell adhesion protein *D*E-cadherin and another through the signaling molecule decapentaplegic.

## Introduction

Adult *Drosophila* structures are derived from clusters of imaginal cells that are specified and determined in the embryo. There are two types of imaginal cells, the imaginal discs and the histoblasts. Histoblasts are nests of larval abdominal cells that are precursors of the fly abdomen, and do not proliferate until metamorphosis. The imaginal discs arise from imaginal cells that invaginate from the embryonic epithelium and become two layered epithelial sacks during larval development. The pattern of these structures is molecularly defined early in development. Imaginal discs proliferate extensively during larval development (Sprey, T.E., Oldenhave, 1974; Haynie and Bryant, 1986). During metamorphosis, imaginal discs evert and differentiate to form the adult appendages and head and thorax epidermis. Imaginal discs are formed by two epithelial layers: the lower disc proper (DP), a columnar epithelium that develops into the principal adult structures, and the upper peripodial squamous epithelium (also known as peripodial membrane) that gives rise mostly to the integumentary cuticle of the adult body wall, being specially important where adjacent discs will suture (Agnès, Suzanne and Noselli, 1999). It has been shown, at least in wing discs, the PE is an indispensable developmental field that has different genetic requirements from wing’s and notum’s fields and it is defined early in development (Baena-Lopez, 2003). The role of the PE has been studied mainly in leg, wing and eye imaginal discs, however, its role in the development of the antenna and proboscis is little known. In the discs where it has been studied, the PE is needed for disc eversion and to control cell proliferation by sending signals to the DP (Milner, Bleasby and Pyott, 1983; Milner, Bleasby and Kelly, 1984). The PE is also important for the establishment and maintenance of disc developmental compartments through direct contact between apposed cells or by transluminal extensions from the PE to the DP (Cho *et al.*, 2000; Gibson and Schubiger, 2000). It has also been shown that the PE secretes exoskeleton cuticle (Milner, Bleasby and Pyott, 1983; Mikołajczyk *et al.*, 1995). The PE has molecularly defined, lineage restricted compartments and it contributes with cells to the DP. In some imaginal discs, such as the wing and leg, cuboidal cells, that are at the border between the PE and the DP are derived from both of these epithelia, while in the eye they are derived only from the PE. This difference in cell lineage contribution makes each disc and its development different, thus stressing the fundamental developmental principle that underlying cellular and molecular processes are commonly used but tissue specific peculiarities define the final morphology or identity of each structure.

Several head structures are directly derived from the eye-antennal lateral PE: the second antennal segment, the vibrissae, the maxillary palps, and the gena (the part of the head to which the jaws are attached) (Bessa and Casares, 2005; Stultz *et al.*, 2006; Lee, Stultz and Hursh, 2007). During early larval development, the major signaling molecules controlling development of the eye-antennal disc are Hedgehog (Hh), Decapentaplegic (Dpp), and Wingless (Wg) which are expressed in the PE but not in the DP. In third instar larvae, the expression of these proteins changes and becomes restricted to the DP (Cho *et al.*, 2000). The mechanisms that establish the dynamic and asymmetric patterns of these signaling molecules are yet to be described but it is known that the asymmetric expression of Hh, Wg, and Dpp in the PE during the second instar refines Delta (Dl) and Serrate (Ser) expression patterns establishing the disc’s dorso/ventral boundary (Cho *et al.*, 2000). Alterations in Dpp regulation affect the development of ventral head structures as it was revealed by the identification the *Dpp*^*s-hc*^ allele which has a mutation in its 5’ eye-antennal enhancer (Stultz *et al.*, 2006). This evidence clearly shows that the interactions between the PE and the DP are critical for the correct patterning and development of the adult structures that emerge during metamorphosis.

The gene *escargot* (*esg*) codes for a transcription factor of the *snail* family. It has five Zn^2+^-finger DNA-binding domains that binds to its consensus sequence 5′-A/GCAGGTG-3′ and two P-DLS-R/K motives that recruit the dCtBP co-repressor, therefore it can activate or repress gene expression depending on the cellular context (Ashraf *et al.*, 1999; Ashraf and Ip, 2001; Cai, Chia and Yang, 2001). It is expressed in all imaginal discs and histoblasts where it has been shown that it is needed for maintaining diploidy and stemness. Esg loss of function in histoblasts promotes premature cell differentiation affecting cuticle development (Hayashi *et al.*, 1993; Fuse, Hirose and Hayashi, 1994). Esg positively regulates *D*E-cadherin expression mediating cell adhesion, motility, survival and differentiation (Tanaka-Matakatsu, Tadashi Uemura, *et al.*, 1996). Depending on their nature, *esg* alleles have a variety of phenotypes that include deletions of tergites and sternites, held out wings, twisted legs, reduced eyes, and null mutants are homozygous lethal.

We have previously reported and characterized a P{GaWB} insertion line (*esg*^*L4*^) that knocks-out *esg* expression. As expected, *esg*^*L4*^ is homozygous lethal while the heterozygous mutant has half of the normal gene dosage. Its GAL4 expression pattern is consistent with the *esg* expression pattern reported in previous literature. This driver expresses GAL4 in the embryonic ectoderm and histoblasts; larval leg, wing and eye-antennal imaginal discs, brain, spiracles and tracheae; adult intestinal stem cells, malpighian tubules, testis and salivary glands. Importantly, we found that *esg* is also expressed in the labial imaginal disc (Sanchez-Díaz *et al.*, 2015).

In the present work we found that lowering the dosage of Esg using RNAi against *esg* (*esg-*RNAi) driven either by *esg*^*L4*^ or by a PE-specific driver, causes abnormalities mainly in the adult antennae maxillary palps and proboscis. These abnormalities are dose dependent; their range of phenotypes goes from the complete loss of the proboscis and antenna to structure duplications of antennae or maxillary palps. We demonstrate that *esg* is expressed in the PE of the antennal disc, and its expression domain also includes the epithelium of the base of the labial disc which is likely to be part of the precursors of the gena. Correct *esg* expression is necessary for the normal development of the antennal and labial discs but its function is different in each of these two discs. When Esg dosage is reduced in the antennae disc, the PE expands over the DP, *D*E-cadherin is reduced and the expression domains of *cubitus interruptus* (*ci*) and *engrailed* (*en*) are affected causing structure malformations of antennae and maxillary palps, however, lowering *dpp* in the expression domain of *esg* does not affect these structures. On the other hand, in the labial discs, a complete loss of the proboscis is caused by lowering *esg* or *dpp* in *esg* expressing cells.

## Results

### *esg* loss of function induces defective development of antennae and proboscis

Since head structures are derived from the eye-antennal and the labial discs, and there is strong *esg* expression in these discs, we decided to investigate the role *esg* in their development. To lower *esg* expression levels we used four different RNAis against *esg* driven by the *esg*^*L4*^ which has a *P{GawB*} insertion at the 5’ of the *esg* gene. The *esg*^*L4*^*>esg-RNAi(v9794)* genotype showed a variety of defects in the development of head structures that included: proboscis loss, where only the labrum can be observed (**Fig. 1A**), antenna duplication (**Fig. 1B**), antennae loss (**Fig. 1C**), and maxillary palp duplication and reduction(**Fig. 1D**). The map of the *esg*^*L4*^ insertion and the location of target sites of the RNAis used are shown in **Figure 1H**. The homozygous *esg*^*L4*^ line is lethal while the heterozygous expresses half the normal *esg* dosage being viable and fertile with no defects in external head structures (**Fig. S1B)**. The other RNAi lines used induced similar phenotypes with varying degrees of penetrance **(Table 1)** that go from subtle proboscis defects, as in *esg*^*L4*^*>esg-RNAi(V28514)* (**Fig. S1C**), to a lethal phenotype, *esg*^*L4*^*>esg-RNAi(34063)*, where only few individuals developed to pupal stages so they had to be analyzed by dissecting them out of the pupal case (**Fig. S1F**). Although *esg* is expressed in all imaginal discs, we found that the most affected structures were the head chemosensory organs: the antennae and the maxillary palps, that are olfactory, and the proboscis that is gustatory. In several of the RNAi expressing lines there was a necrotic area in the proboscis area. These organisms are able to eclose but they emerge without proboscis and the necrotic region remains attached to the pupal case (**Fig. 1F**).

**Table 1.** Phenotypes observed when RNAis against esg are driven by the drivers *esg*^L4^> or C311>

| ***esg*<sup>L4</sup>>** | **V/L** | **Larval phenotype** | **Adult phenotype** | **C311 >** | **V/L** | **Larval phenotype** | **Adult phenotype** |
| --- | --- | --- | --- | --- | --- | --- | --- |
| **28514** | **V** | Normal | Partial proboscis defects, antennae and maxillary palps intact | **V28514** | **V** | Not tested | Not tested |
| **V9794** | **V** | Third instar imaginal discs are partially or completely developed | Damage in the antennae and maxillary palps and partial damage or complete loss of proboscis | **V9794** | **V** | Normal | Damage to antennae. |
| **V9793** | **L** | Third instar imaginal discs are sometimes partially developed | Damage to antennae and maxillary palps and complete proboscis loss in all animals | **V9793** | **L** | Third instar larvae imaginal discs are all under developed or missing | Pupate but no imago |
| **34063** | **L** | Third instar imaginal discs are all under-developed or missing | Pupate but do not eclose, dissected pharates always have complete proboscis loss and damage to antennae and maxillary palps. | **34063** | **L** | Third instar larvae imaginal discs are all under developed or missing | Pupate but no imago |

The first column of each side of the table shows drivers *esg*<sup>L4</sup>> or c111> and the different RNAis used. V = viable and L = lethal

**Fig. 1.**
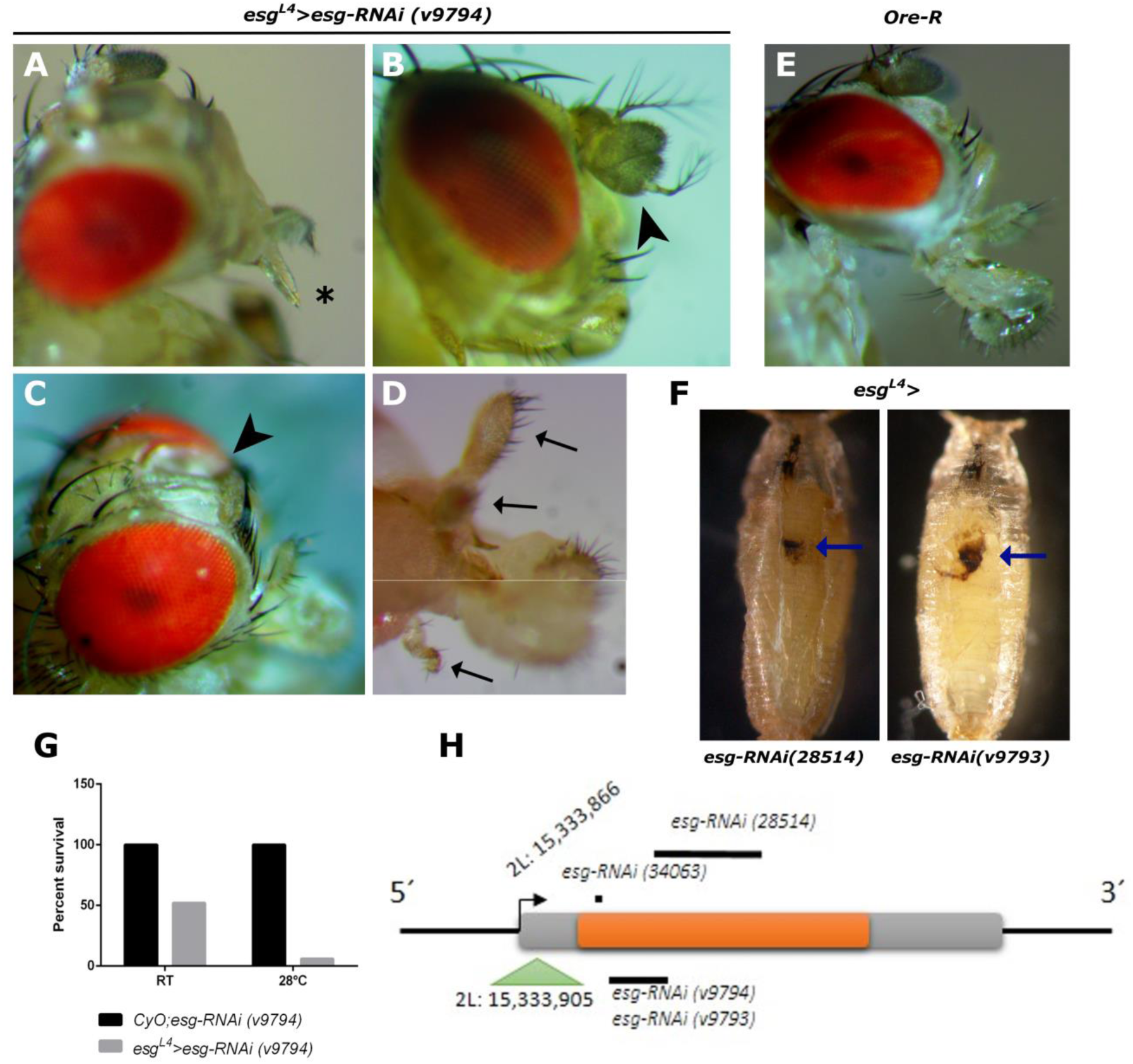
Range of chemosensory organs defects in flies expressing *esg-RNAi(v9794)*. **(A-D)** Different individuals showing the diversity of phenotypes found in *esg*^*L4*^*>esg-RNAi(v9794)*. **A)** Proboscis loss (asterisk). **B)** Antennae duplication (arrowhead). **C)** Antennae loss (arrowhead). **D)** Extreme phenotype where one individual has a duplication of the maxillary palp on one side and reduction of the maxillary palp on the other side (arrows), the panel is divided because it was impossible to focus both sides simultaneously. **E)** Normal chemosensory organs of a wt fly (Ore-R). **F)** Pupae of the corresponding genotypes where necrotic cells around the proboscis area can be seen (arrows). **G)** Viability as a function of temperature. **H)** Schematic representation of *esg* genomic region showing the insertion site of the P{GaWB} element (*esg*^L4^) and the location of the target sites of the RNAis used.

*esg*^*L4*^*>esg-RNAi(v9794)* genotype has the whole range of abnormalities that appeared with all the other RNAi lines while being fertile and viable, although its viability was severely reduced when incubated at 28°C degrees when compared with development at room temperature (**1G**). We used this line to calculate the frequency of different phenotypes found, but as there was a wide range of phenotypes, we classified them by the affected organ according to their function (gustatory, olfactory or both). At room temperature, we found that 72% of individuals had defects on the proboscis (gustatory); 10% had defects on the antennae and/or maxillary palps (olfactory); and 18% had defects on both, proboscis and maxillary palps and/or antennae. In this genotype there were no individuals that had normal chemosensory organs. At 28°C, there were only 8 individuals that survived to adulthood, 6 of them had gustatory defects and 2 with both gustatory and olfactory defects **(Table 2)**. Quantitative PCR of eye-antenna discs showed that reduced *esg* expression levels inversely correlate with the severity of the phenotypes observed and in the over expressing line (*esg*^*L4*^*>P{EP}esg*), normal esg levels are recovered (**Fig. S2**).

**Table 2:** Phenotypes observed at different temperatures

| Phenotype | RT<br>*esg<sup>L4</sup>>esg-RNAi (v9794)*<br>(%) | 28°C<br>*esg<sup>L4</sup>>esg-RNAi (v9794)*<br>(%) |
| --- | --- | --- |
| Gustatory | 55 (72) | 6 (75) |
| Olfactory | 8 (10) | 0 |
| Both | 14(18) | 2 (25) |
| Normal | 0 | 0 |

### Defects induced by *esg* RNAi on head chemosensory organs are not caused by an increase in cell death

Mutations in *dpp* cause similar defects in the maxillary palps as the ones observed in this work. Loss of *dpp* in the PE induces apoptosis in the DP (Stultz *et al.*, 2006). In order to find out if the defects we observed in the *esg* hypomorphic mutant lines were caused by increased cell death during development, we performed TUNEL assay on third instar larvae eye-antenna and labial imaginal discs (**Fig. 2**). In labial discs of wt and *esg*^*L4*^ and *esg*^*L4*^*>esg-RNAi(v9794)* lines there was no significant differences in the number of TUNEL positive cells. On the eye-antenna disc, there were more dying cells in the *esg*^*L4*^ and *esg*^*L4*^*>esg-RNAi(v9794)* in comparison to the control line, however there was no significant difference between *esg*^*L4*^ and *esg*^*L4*^*>esg-RNAi(v9794)*. As the latter always shows some type of head structure defects but *esg*^*L4*^ never presents this type of malformations, we concluded that it is not likely that cell death accounts for the phenotypes observed.

**Fig. 2.**
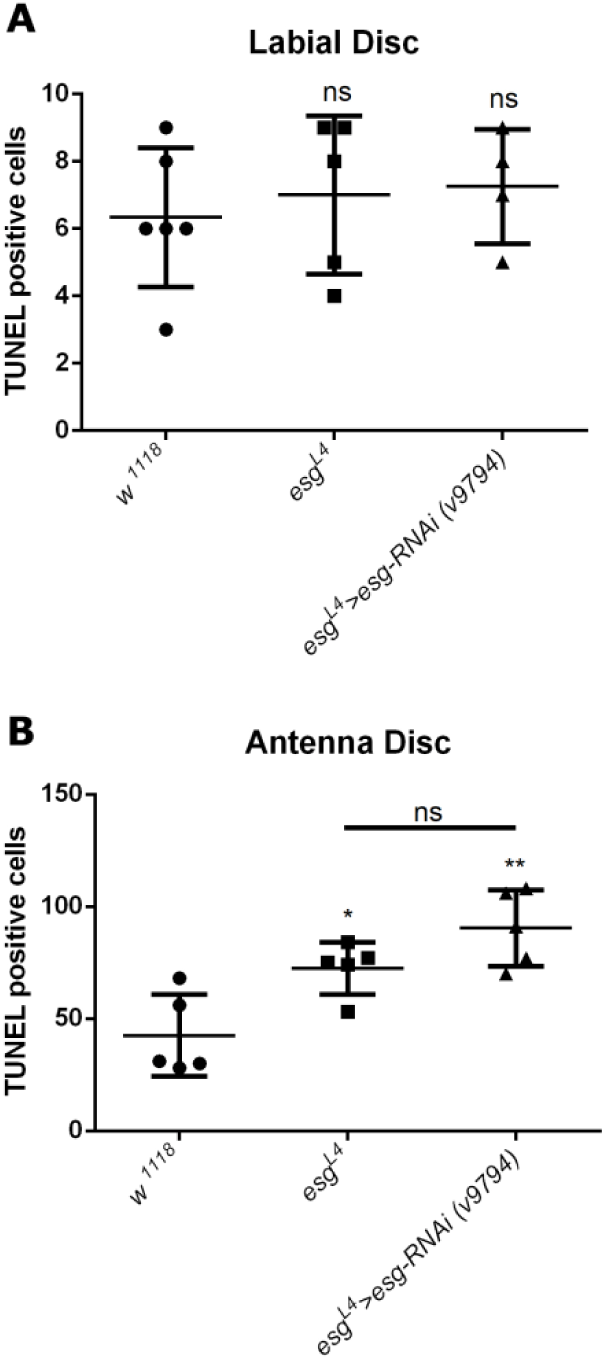
Cell death is not the cause of head structures defects. Number of TUNEL positive cells in labial **(A)** and antennal discs **(B)**.

### *esg* defines a morphogenetic domain within the peripodial epithelium connecting the labial and antennal discs

Given the strong phenotypes that we find in head structures when *esg*-RNAis are expressed, we decided to characterize how the expression domain of *esg*^*L4*^ changes as function of its dosage. In order to do this, we rescued the endogenous *esg* levels using *P{EP}esg* to compare it with *esg*^*L4*^ that has half dosage and with RNAi expressing lines that have even lower levels. We found that the mCD8::GFP expression domain has higher GFP expression and expands whenever any *esg-*RNAi is expressed. In **Fig. 3** we show an example of this phenotype in the *esg*^*L4*^*>esg-RNAi(v34063)/UAS-mCD8::GFP* line. When the UAS-*esg*-RNAi is expressed together with UAS-mCD8::GFP using *esg*^*L4*^ as a driver, the number of cells expressing GFP in the embryo augments (**Fig. 3 A** and **B)**. In labial discs, when the Esg levels are rescued using *esg*^*L4*^*>P{EP}esg* the region that expresses GFP is restricted to the base of the disc and perimandibular cells (**Fig. 3 C**). In *esg*^*L4*^ labial imaginal discs, GFP expression is still restricted the base of the disc and the perimandibular region, but there is a stronger signal in the perimandibular cells (**Fig 3 D**). In *esg*^*L4*^*>esg-RNAi(v34063)/UAS-mCD8::GFP* where the Esg levels are even lower than in *esg*^*L4*^*>UAS-mCD8::GFP*, the GFP expression of the perimandibular cells becomes stronger, the labial disc is unrecognizable, and in its place a large cluster of GFP positive cells is found (**Fig. 3 E)**. This large cluster of cells perimandibular cells are likely to be the necrotic cells found later in development attached to the pupal case.

**Fig. 3.**
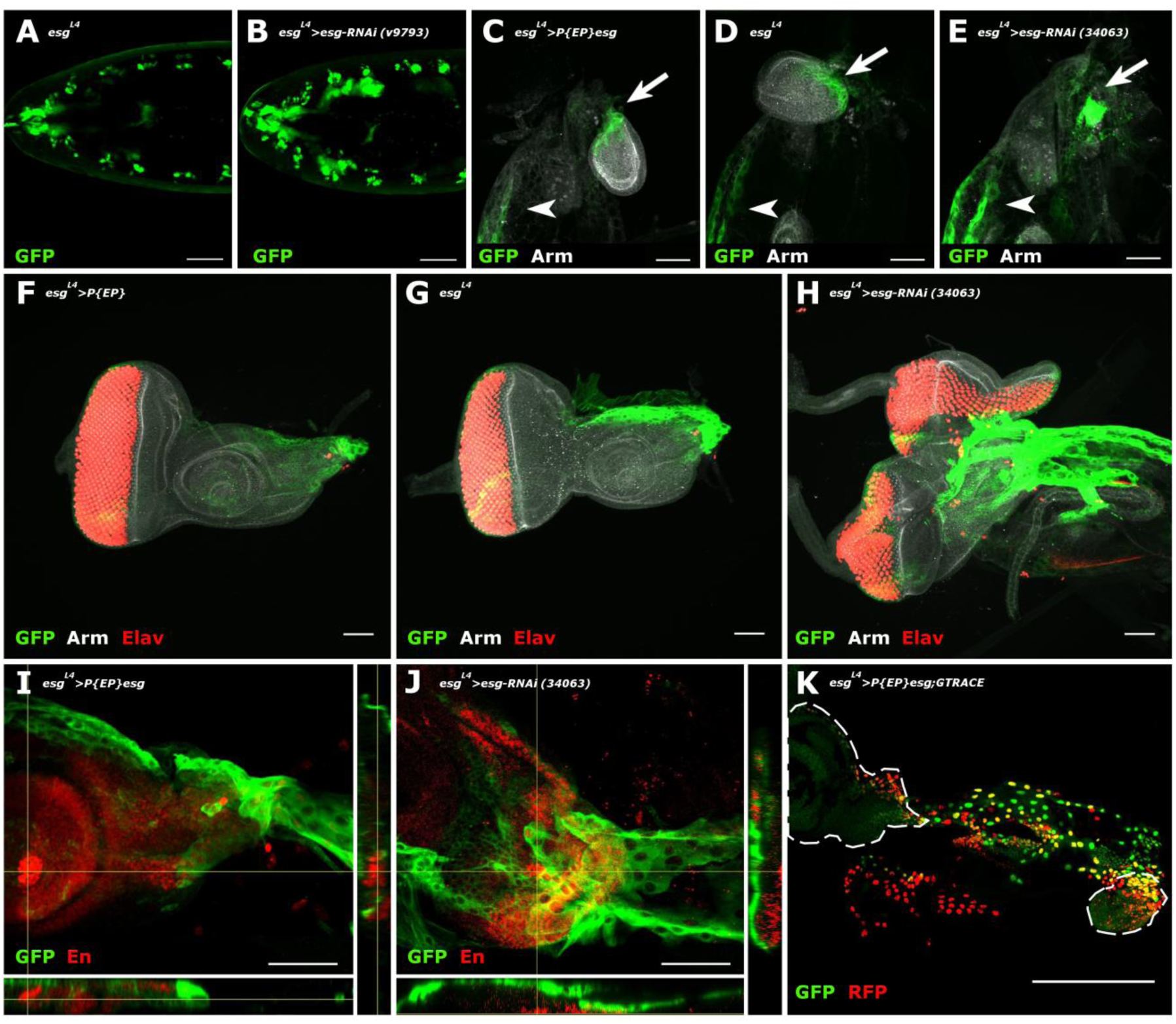
*esg* defines a morphogenetic domain within the peripodial epithelium connecting the labial and antennal discs. GFP expression driven by *esg* in lines with different level of *esg* expression. **A)** Embryonic expression pattern of *esg*^*L4*^. **B)** Embryonic expression pattern of *esg*^*L4*^ with an *esg-*RNAi to lower *esg* levels induce more GFP expressing cells. **C-E)** Third instar larvae mandibular region of the labeled genotypes. GFP signal increases with decreasing levels of *esg* expression. **C)** In the *esg*^*L4*^*>P{EP}esg/UAS-mCD8::GFP* line, GFP positive cells are restricted to the base of the labial disc (arrow) and the perimandibular cells (arrowhead). **D)** In *esg*^*L4*^*/UAS-mCD8::GFP* an increase in the area covered by GFP positive perimandibular cells (arrowhead) at the labial disc (arrow) can be observed. **E)** In *esg*^*L4*^*>esg-RNAi(v34063)/UAS-mCD8::GFP)* perimandibular cells (arrowhead) are bigger and cover a greater area, the labial disc is completely lost and in its place a large cluster of GFP positive cells (arrow) is found. **F-H)** Eye-antenna discs of the labeled genotypes. **F)** In the *esg*^*L4*^*>P{EP}esg/UAS-mCD8::GFP* line rescued dosage of *esg* expression restricts GFP to the base of the disc and the PE. **G)** Half dosage of *esg* (*esg*^*L4*^*/UAS-mCD8::GFP*) causes an expansion of the area where GFP is expressed. **H)** Lower levels of *esg* dosage (*esg*^*L4*^*>esg-RNAi(v34063)/UAS-mCD8::GFP)* extends the GFP expression domain covering and invading the dorsal area of the antennal disc. Cellular markers in **A** to **H** have the following channels: green = GFP, grayscale = β-catenin and red = pan neural marker Elav. **I-J)** Higher magnification of the peripodial membrane of the corresponding genotypes. **I)** Rescued *esg* organisms (*esg*^*L4*^*>P{EP}esg/UAS-mCD8::GFP*) show GFP expression that is restricted to the peripodial epithelium of the antennal disc. **J)** GFP positive cells of an antenna disc with very low *esg* dosage (*esg*^*L4*^*>esg-RNAi(v34063)/UAS-mCD8::GFP*), the antennal disc stalk is lost, the perimandibular cells invade the posterior part of the antennal disc. green = GFP, red = Engrailed. **K)** Expression of the GTRACE marker in the perimandibular region: *esg* expressing cells are a continuous field that connects the labial with the antennal disc. In this panel, cells with active *esg* expression are marked with RFP; cells that expressed *esg* but are no longer expressing are marked with GFP; yellow cells express both RFP and GFP. Antennal and labial discs are delimited by the dashed lines. All scale bars = 50 µm except K scale bar = 200 µm

In antennal discs, with normal levels of *esg* (*esg*^*L4*^*>P{EP}esg*), most of the GFP expression is concentrated in the stem of the antennal disc, with little expression on the dorsoposterior side (**Fig. 3F**). With half a dosage of *esg* the dorsoposterior domain expands covering the whole dorsal rim of the antennal disc (**Fig. 3 G**). When *esg* levels are lowered using RNAi, the GFP expressing cells expand even more covering most of the antennal disc (**Fig. 3 H**). Importantly, the cells expressing *esg* are restricted to the PE of the third instar eye-antenna disc and not to the DP; this can be observed in the transversal sections depicted in **Fig. 3 I and J** were the Engrailed (En) signal marks the DP.

To define the domains where *esg* has been expressed we used the developmental marker GTRACE (Evans *et al.*, 2009) where *esg*^*L4*^ drives RFP and flipase which in turn activates permanently the expression of GFP by removing a stop codon that interferes with its expression after *esg* is no longer expressed. The *esg*^*L4*^*>P{EP}esg:UAS-GTRACE* line showed that *esg* was expressed early in both the antennae and labial DP but is no longer expressed in these tissues at this stage as they are marked in green. However, the perimandibular cells and the antennal PE are expressing *esg* at this stage, as can be observed by the red and yellow signal. Interestingly, the peripodial expression of *esg* is a continuum that connects both discs (**Fig. 3K)**.

### Defects caused by loss of *esg* in the antennal disc involve the peripodial epithelium

In order to demonstrate that the phenotype observed is caused by *esg* loss of function in the PE, we used a PE specific driver (*C311-GAL4*) to knockdown *esg* expression. We verified the expression pattern of this driver using UAS-GFP and compared it with *esg*^*L4*^ driver. Effectively, C311-GAL4 marks the whole PE of the eye-antennal disc while *esg*^*L4*^ only marks a subdomain of this PE (**Fig. 4 A and B**). Expression of an *esg-RNAi* under C311 control reduces the antennal disc in a similar way as the expression of this RNAi under the control of *esg*^*L4*^ does (**Fig. 4 C and D**).

**Fig. 4.**
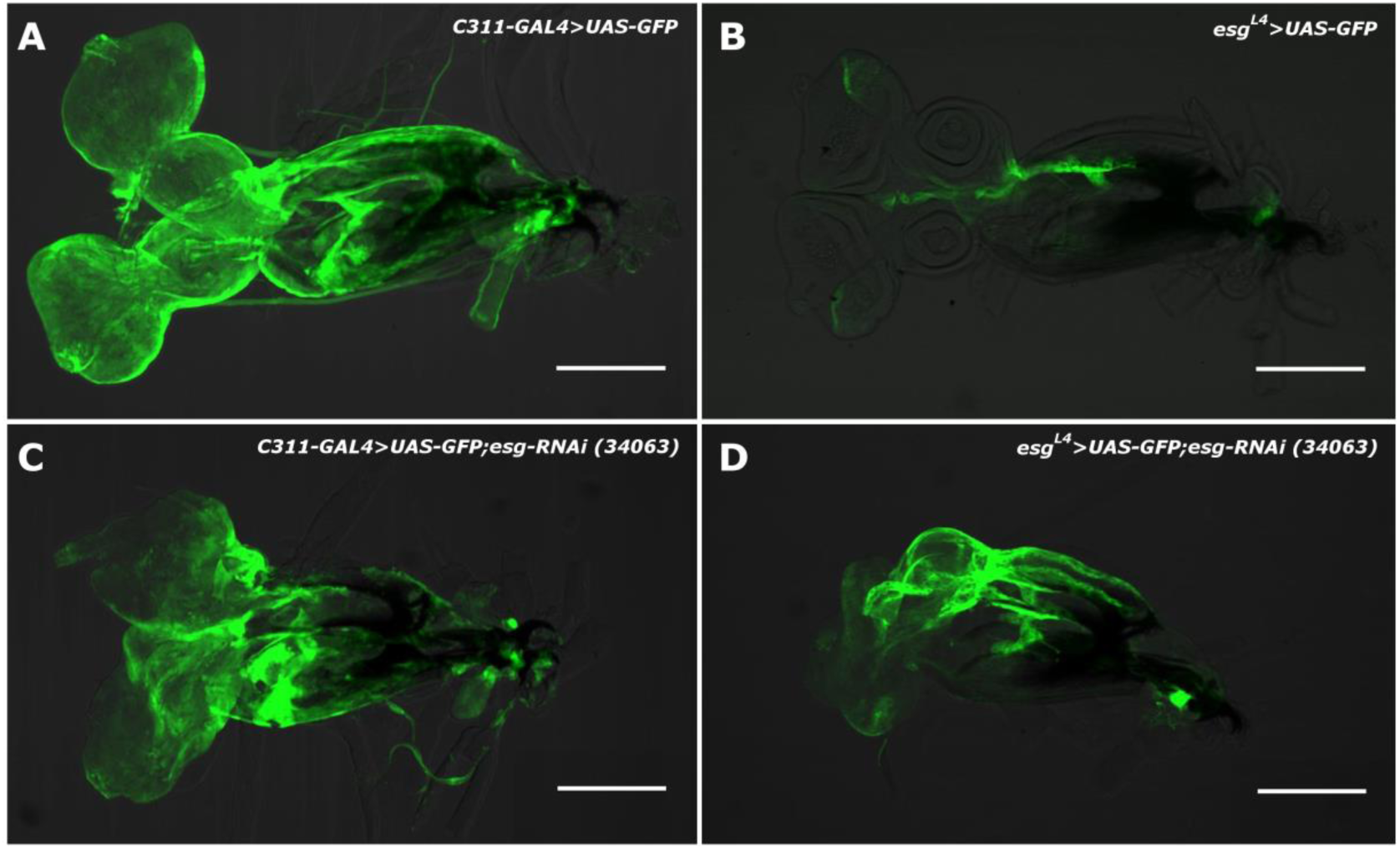
Expression pattern of the C311-GAL4 driver. **A)** Expression of GFP driven by C311-GAL4 marks the PE of the eye-antennal discs, the perimandibular cells and the PE of the labial discs. **B)** Expression of GFP driven by *esg*^*L4*^ marks a subdomain of the C311 expression pattern. **C)** Knockdown of *esg* in C311 pattern only affects the antennal discs, while leaving intact the eye and labial discs in a similar way as *esg*^*L4*^ does **(D)**. Scale bar = 200 µm

Analyzing the antenna discs in higher magnification and using as anterior marker Ci and En as posterior marker we found that in the *C311-GAL4>UAS-GFP* antennal discs the anterio-posterior domain is well defined and the expression of GFP is in the PE and in the stalk of the antennal disc **(Fig 5 A)**. Adult organisms are normal (**Fig 5 B**). Knocking down *esg* expression with this driver induced a similar phenotype to the one observed when *esg-*RNAis were driven by *esg*^*L4*^. Expression of *esg-RNAi(v34063)* had a pupal lethal phenotype. Third instar larval antennal discs from these animals are reduced, and a similar expansion of the PE to the one observed when this RNAi is driven by *esg*^*L4*^ is observed (**Fig. 5C**). When the relatively mild *UAS-esg-RNAi(v9794)* was used, we found malformations in the antenna **(Fig. 5D**). The resulting phenotypes observed when driving different *esg*-RNAs using the PE specific driver C311-GAL4 are summarized in **Table 1**. These results suggest that *esg* is expressed in the PE of the antenna disc, and that it restricts the expansion of the PE over the DP. Relevantly, the expression *of esg-RNAi* with this PE driver did not have any observable phenotype in the proboscis.

**Fig. 5.**
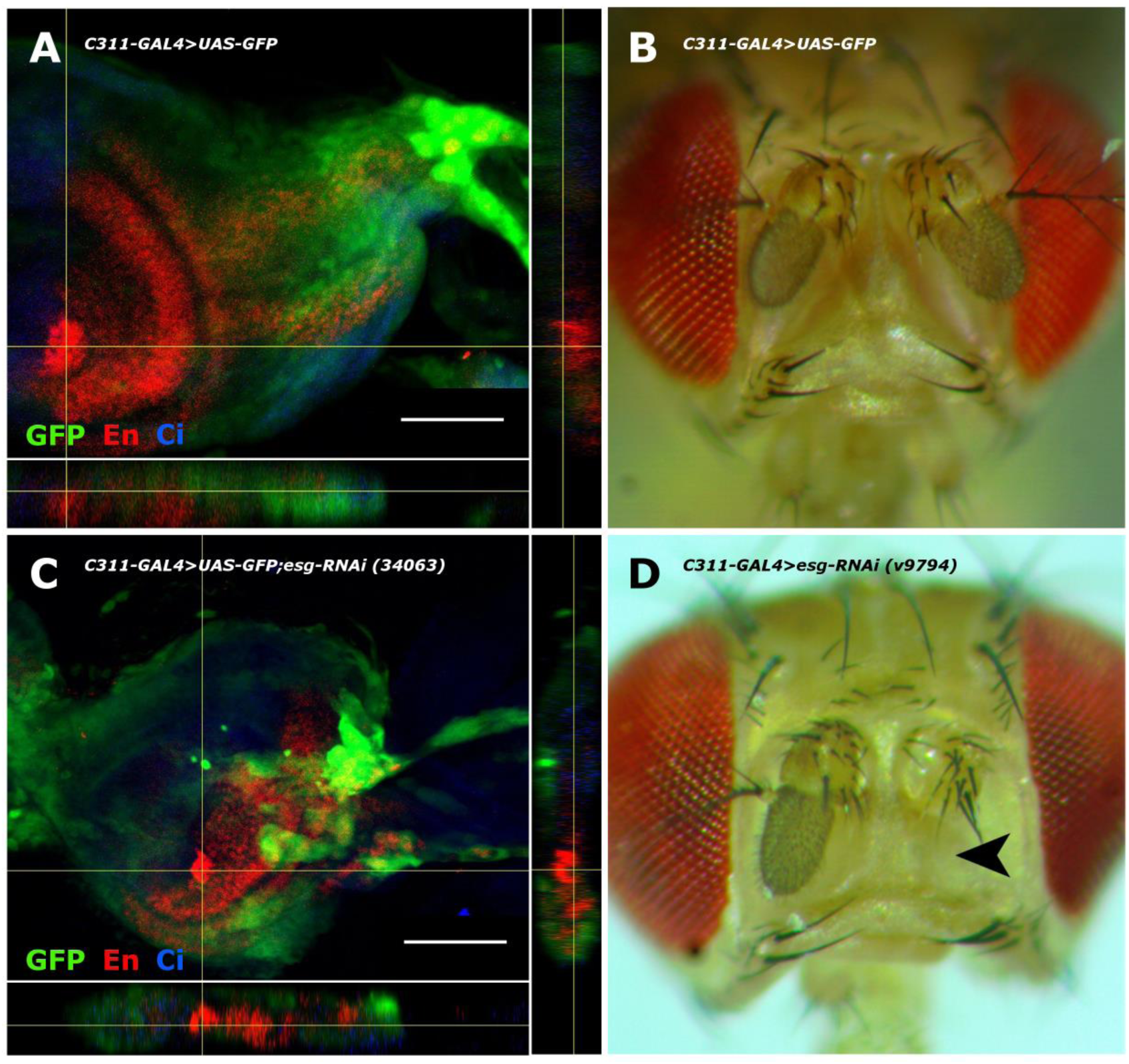
Specific *esg* knockdown in the peripodial epitelium produces head structures defects and expansion of the peripodial epithelium. **A)** Expression of GFP driven by C311-GAL4. GFP is only expressed in the PE as it can be observed in transversal section insert. **B**) C311-GAL4>GFP has a wt adult phenotype. **C)** The PE expands over the DP when the *esg-RNAi(v34063)* is driven by C311; this genotype dies at pupal stages. **D)** Adult phenotype observed when the *UAS-esg-RNAi(v9794)* is driven by C311-GAL4, loss of the distal part of one of the antennae can be observed (arrowhead). Channels: Green = GFP, Red = Engrailed, Blue = Cubitus interruptus. Scale bar = 50 µm

### *esg* loss of function causes a reduction in *D*E-cadherin in antennal imaginal discs

It has been shown that during tracheal system development, *esg* is a positive regulator of *D*E-cadherin (Tanaka-Matakatsu, Tadashi Uemura, *et al.*, 1996). To determine if the downregulation of *D*E-cadherin could be participating in the phenotypes observed when *esg-RNAis* are driven by *esg*^*L4*^, we performed western blot analysis of *esg*^*L4*^*>esg-RNAi(v34063)* third instar larvae (**Fig. 6 A and B)** and found that that *D*E-cadherin expression is strongly reduced, thus explaining the lethality observed in this genotype. Conversely *D*E-cadherin levels are rescued when the P{EP}*esg* is expressed. We immunostained eye-antennal and labial disc with anti-*D*E-cadherin and found that in *esg*^*L4*^ *D*E-cadherin there is a clear reduction in the periphery of the antennal segments while the effect is less obvious in the eye disc (**Fig. 6D**). When *esg* levels are further reduced in the line *esg*^*L4*^*>esg-RNAi(v34063)* there is an even stronger reduction of *D*E-cadherin in the antennal disc, with clear alterations in the disc morphology. *D*E-cadherin pattern becomes de-localized, the segments cannot be identified anymore and invasion of the perimandibular cells can be observed on the dorsal side of the disc **(Fig 6E)**. On the other hand, the *D*E-cadherin levels in the eye disc do not seem to be downregulated with the lowering dosages of Esg. This is also the case of the labial discs where *D*E-cadherin levels do not seem to be downregulated. In *esg*^*L4*^ *D*E-cadherin levels appear to be very similar to the levels of the control *w*^*1118*^ **(Fig. 6 D and E)**. In the *esg*^*L4*^*>esg-RNAi(28514)* line, the size of the disc is reduced, but the levels of *D*E-cadherin appear to be maintained throughout the disc. When *esg* levels are further lowered in *esg*^*L4*^*>esg-RNAi(v34063)*, the disc completely disappears and the tissue becomes extremely fragile (**Fig. 6G**). This suggests that *esg* is downregulating *D*E-cadherin specifically in the antennal disc but not in the labial or eye disc. Therefore, the phenotypes of loss of proboscis and damage to the antenna and maxillary palps seem to be caused by two different pathways.

**Fig. 6.**
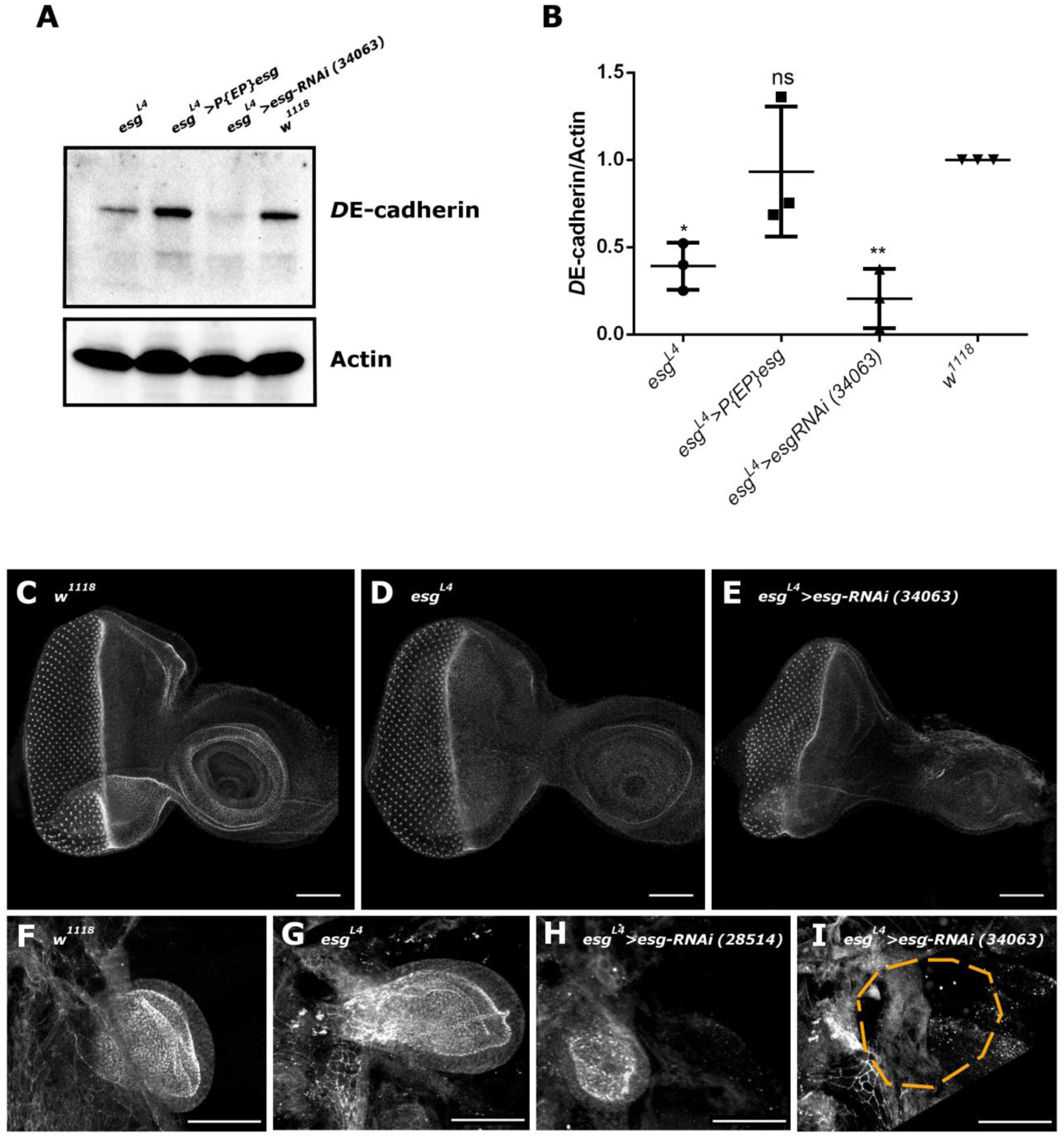
Esg positively regulates *D*E-cadherin. **A)** Western-blot of the corresponding genotypes. *D*E-cadherin expression is lower in *esg*^*L4*^ and practically disappears when the RNAi is expressed. **B)** Densitometric quantitation of western blot in **H** (n = 3). All genotypes are compared to *w*^*1118*^ **(C-E)** *D*E-cadherin expression in eye-antenna imaginal disc. **C)** Control (*w*^*1118*^) *D*E-cadherin is clearly expressed in the periphery of the precursors of the antennal segments. **D)** *esg*^*L4*^ shows a slight decrease in the expression of *D*E-cadherin in the precursors of the antennal segments. **E)** In *esg*^*L4*^*>esg-RNAi(34063) D*E-cadherin expression is severely reduced and the eye-antenna imaginal disc is deformed. **F-I)** *D*E-cadherin expression in the labial imaginal disc of the corresponding genotypes. **F)** Control (*w*^*1118*^), the labial discs have a characteristic expression of *D*E-cadherin. **G)** In *esg*^*L4*^, the labial disc has very similar *D*E-cadherin expression. **H)** The size of the labial disc is greatly reduced when the RNAi *esg-(281514)* is expressed; *D*E-cadherin can still be detected in its corresponding domains. **G)** When *esg-RNAi(34063)* is expressed, labial imaginal disc are completely loss and the tissue becomes very fragile. Scale bar = 50 µm

### The anteroposterior axis of the antennal disc becomes disorganized when there is *esg* loss of function

Low *esg* dosage induces PE expansion in the eye-antenna disc. There was no observable expansion in the leg and wing discs and in our hands we did not observed malformations in these organs. Since the PE is an important source of morphogenetic signals and the PE expansion correlates with developmental defects in the antennae and maxillary palps, we decided to observe if there was an alteration of the anteroposterior axis of the antennal disc. To label the anterior compartment we used an antibody against *Cubitus interruptus* (Ci) and for the posterior compartment we used an antibody against Engrailed (En) (**Fig. 7**). In the heterozygous *esg*^*L4*^ background that shows no external defects, the anteroposterior compartments of the antennal disc are as well defined as in *w*^*1118*^(**Fig. 7 A** and **I**); this can also be observed in the wing disc (**Fig. 7 E**). When the *esg-RNAis* are expressed, the anteroposterior compartments of the wing discs remain well defined (**Fig. 7 F** and **G**) whereas in the antennal disc the anteroposterior domains become disorganized (**Fig. 7 B** and **C**). The disorganization of these compartments was variable and this is consistent with the phenotypical diversity that we observed in adult individuals of these genotypes. In some extreme cases we found duplicated *en* domains that may explain the apparition of duplicated antenna in adults (**Fig. 7 K** and **L**). Importantly when *esg* gene dosage is restored in a *esg*^*L4*^*>P{EP}esg* the organization of the anteroposterior compartment is restored in the antennal disc (**Fig. 7 D)** and there is still no alteration in the wing disc **(Fig. 7 H)**.

**Fig. 7.**
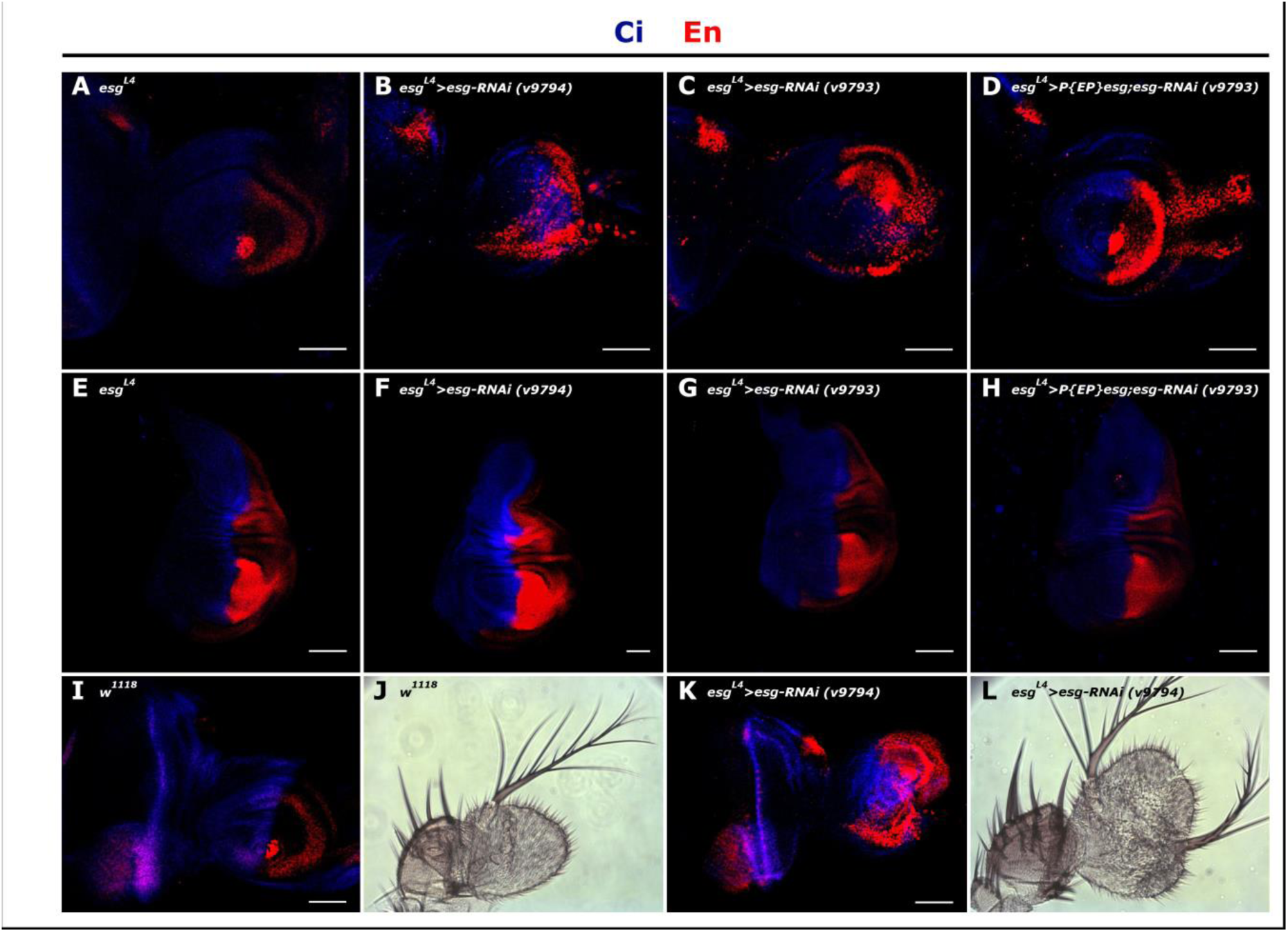
The anteroposterior (AP) axis is distorted when *esg* is knocked down. Immunolocalization of posterior domain marker Engrailed (red), and anterior domain marker Cubitus interruptus (blue), in third instar larvae antenna discs (upper panels) and wing discs (middle panels) of the corresponding genotypes. Half *esg* dosage (*esg*^*L4*^) does not have an apparent adult phenotype and the AP domains are well defined in both antenna and wing discs (**A** and **E**). When the *esg-RNAis* are used (v9794 and v9793) adult head structures are consistently affected and the AP domains become disorganized only in the antenna discs (**B** and **C**) and not in the wing discs (**F** and **G**). Restoring *esg* dosage with a *P{EP}esg* while expressing the RNAis rescues the AP compartmentalization (**D**). In the wing discs there is no effect regardless of the genotype (**E-H**). Some individuals presented duplication of the En domain in the antenal discs, which could explain the apparition of duplicated antennae in adults (**I-L**). **I**) Immunolocalization of En and Ci in *w*^*1118*^ eye antenna disc. **J**) Normal *w*^*1118*^ antenna. **K**) Immunolocalization of En and Ci in *esg*^*L4*^*>esg-RNAi(v9794)*. **L)** Duplicated antenna of a *esg*^*L4*^*>esg-RNAi(v9794)* adult. Scale bar = 50 µm

### Decapentaplegic knock down in the peripodial epithelium phenocopies *esg* loss of function in the labial disc but not in the antennal disc

Mutations in the *decapentaplegic* (*dpp*) gene produce duplications or loss of the maxillary palps. *dpp* is critical for imaginal disc compartmentalization, it is expressed in the PE of the eye-antennal disc and *dpp* de-regulation generates defects of adult head structures (Stultz *et al.*, 2006). We expressed a *dpp-RNAi* using the *esg*^*L4*^ driver in order to phenocopy malformations in the maxillary palps. Unexpectedly, palps were not affected in this genotype but there was a total lack of proboscis in a similar way as when *esg-RNAi* are driven by *esg*^*L4*^ (**Fig. 8 D-F**). The labial disc is reduced but interestingly, the anteroposterior axis of the labial disc is not affected **(Fig. 8 B and C)**. The antenna or the maxillary palps are normal and their anteroposterior compartments are not affected either (**Fig. 8 H** and **I, Table 3)**. These results suggest that the defects observed in *esg* mutants are caused by different pathways in gustatory and olfactory organs. Loss of *esg* in the labial disc does not alter the establishment of the anteroposterior axis and the loss of the proboscis depends on *dpp* expression. Lowering of *dpp* in the same tissues where *esg* is expressed causes the loss of the proboscis thus implicating an interaction of *esg* and *dpp* in proboscis development. As the expression domains of *esg* and *dpp* do not correlate at this stage **(Fig. 8 G)**, its interaction must have occurred earlier in development. On the other hand, *esg* loss in the antennal disc alter the establishment of the anteroposterior domain, but this does not happen when *dpp* is lowered in cells expressing *esg*, explaining the lack of phenotype observed in the structures derived from this disc.

**Table 3:** Effect of *dpp* downregulation in the chemosensory organs

| Phenotype | *CyO;dpp-RNAi*<br>(%) | *esg<sup>L4</sup>>dpp-RNAi*<br>(%) |
| --- | --- | --- |
| Gustatory | 0 | 99.5 (222/223) |
| Olfactory | 0 | 0 |
| Both | 0 | 0 |
| Normal | 100 (223/223) | 0.5 (1/223) |

**Fig. 8.**
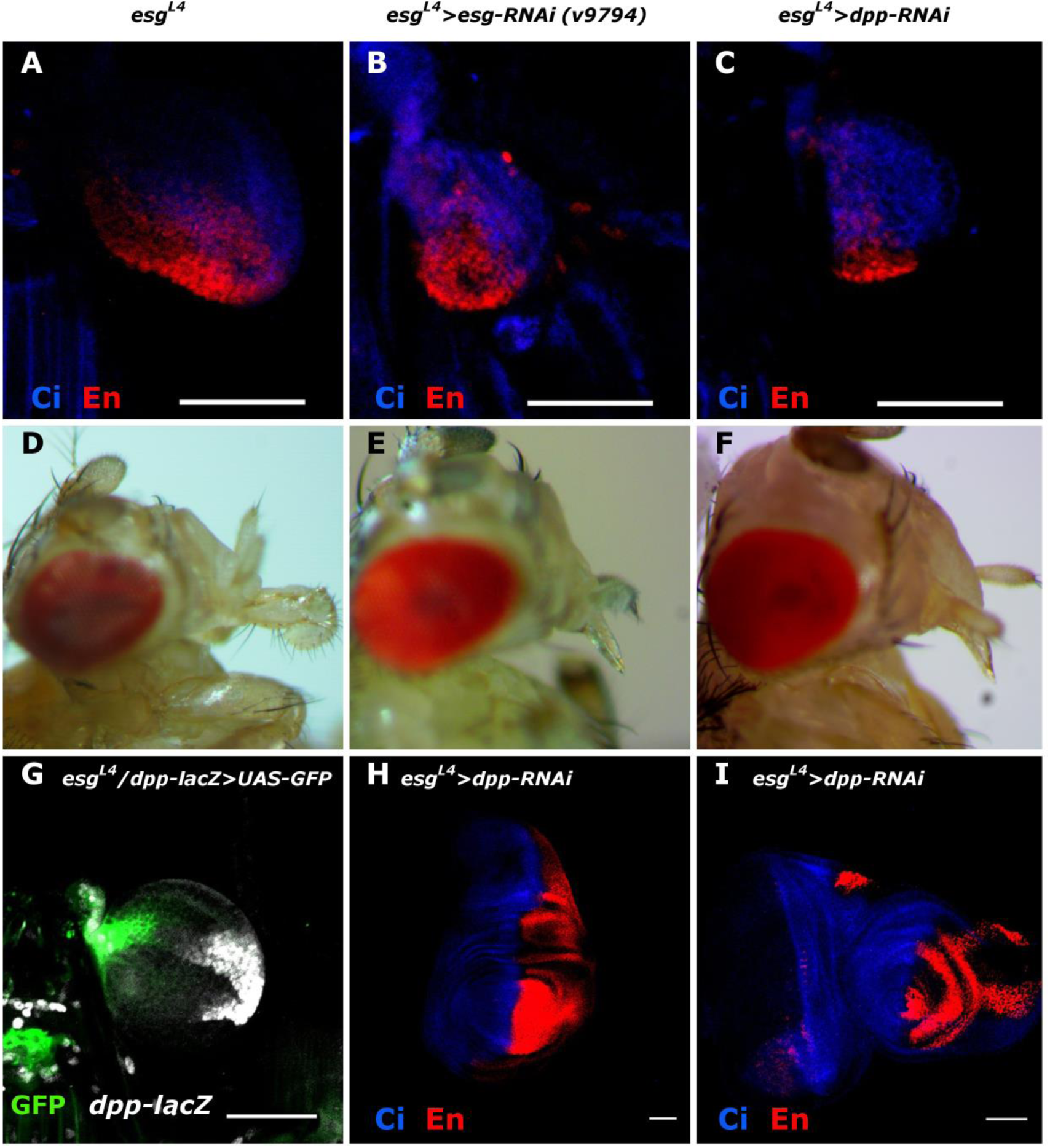
Downregulation of *dpp* in the *esg* domain induces proboscis loss. **A-C)** Labial discs of the corresponding genotypes. There is a reduction in the size of the discs when *esg*^*L4*^ drives *esg-RNAi* **(B)** or *dpp-RNAi* **(C)**, but the anteroposterior compartments are not affected. **D-E)** Adult phenotypes of the corresponding genotypes. There is a complete loss of the proboscis when *esg*^*L4*^ drives *esg-RNAi* **(E)** and when *esg*^*L4*^ drives *dpp-RNAi* **(F). G)** β-Galactosidase (grayscale) and GFP (green) expression in the *esgL4>UAS-GFP; dpp-LacZ* line. Both *dpp* and *esg* are expressed in the labial discs, but their expression domains are complementary. Antenna **(H)** and wing **(I)** discs where the RNAi against *dpp* is driven by *esg*^*L4*^. Neither disc presents alterations in their anteroposterior domains thus explaining the lack of phenotypes in antennae and maxillary palps. Blue = Ci, Red = En. Scale bar = 50 µm

## Discussion

The transcriptional factor *esg* is involved in many important developmental processes that affect a wide variety of tissues and it is clear that cell differentiation and destiny is profoundly influenced by *esg* levels. Different *esg* levels alter tissue development and have a variety of phenotypes. We have previously shown that half genetic dosage of *esg* causes nicotine sensitivity but has no external phenotypes. Further lowering of *esg* dosage has increasing penetrant phenotypes that can be observed in adult structures while null mutants are lethal. In this work we found that *esg* is important in antennal and labial disc development. The developmental defects in the antennae, maxillary palps and proboscis become more penetrant as *esg* dosage diminishes; the phenotype range goes from reduced or malformed structures to missing or duplicated ones. The role of *esg* in the phenotypes observed is confirmed by the fact that they can be suppressed by counter acting the effect of RNAis by recovering expression of this transcription factor using *P{EP}esg.* We found that these malformations are caused by different pathways in the labial and antennal discs. In the labial discs, the pathway is *dpp* dependent and does not alter the anteroposterior compartments. In the antennal disc, the pathway is independent of *dpp*, expression of *esg* is needed in the PE and reduction of *esg* expression causes an expansion of the PE over the DP disorganizing the anteroposterior compartments.

### Esg participation in the antennal disc

It has been shown that *D*E-cadherin is an Esg target. Here we show that this cell adhesion protein becomes specifically down regulated in the antennal disc in genotypes with low Esg levels. The loss of cell adhesion can induce cell death (anoikis) however, our results do not support this possible explanation for the phenotypes found. It is more likely that lack of *D*E-cadherin is altering cell to cell adhesivity allowing them to be more invasive and affecting cell signaling thus modifying cell determination and differentiation of the discs. Previous works have shown that lower levels of *D*E-cadherin have phenotypes with variable expressivity that include: abnormalities in proliferation and morphogenetic movements, loss of epithelial structures and loss of cell polarity(Wang *et al.*, 2004). It is well documented that E-cadherin reduction reduces cell adhesion and promotes tissue invasivity (Conacci-Sorrell *et al.*, 2003). It has also been reported that *esg* regulates cell motility thru the modulation of actin in a manner that is independent of its capacity to regulate cell adhesion (Tanaka-Matakatsu, T Uemura, *et al.*, 1996). The careful analysis of the *esg*^*L4*^ expression pattern suggests that the perimandibular cells and the PE of the antennal disc are a continuous developmental field that physically connects the discs that conform the head and probably serves as a scaffold involved in the growth and differentiation of these discs; the idea that the PE and the perimandibular cells are a continuous developmental field is supported by the fact that the peripodial specific marker and driver C311-GAL4 also directs GFP to this tissue in a pattern that is very similar to the one of *esg*^*L4*^. The expansion of the PE field alters the extension of the molecular signals that it conveys. Our results are consistent with the observations of Gibson and Schubiger that the PE regulates the development of the DP, probably by transluminal signals (Gibson and Schubiger, 2000). As each disc has its own molecular developmental history the interpretation of the ectopic peripodial signal is different and therefore causes disc specific phenotypes. On the other hand, It is reasonable to assume that the expansion of the PE would cause the appearance of supernumerary structures or amorphous tissue, but the developmental evidence shows that this excessive tissue becomes necrotic during metamorphosis as can be observed in **Fig. 1F**, further supporting the idea that the loss of *esg* causes miss-differentiation of this tissue that eventually dies for lack of a proper developmental environment.

### Esg participation in the labial disc

To our knowledge there are no reported mutants that have a complete proboscis loss. One the other hand, one of the few genes whose mutations affect the maxillary palps in a similar way as *esg* does is *dpp*. Interestingly, lowering *dpp* in the *esg*^*L4*^ expression domain causes no effect on the antennae or maxillary palps. The failure to phenocopy *dpp* mutations that affect the maxillary palps when *esg*^*L4*^ drives the expression of the *dpp* RNAi, implies that in these tissues *esg* is probably not expressed in the same regions as *dpp*. However, reduction of *dpp* with *esg*^*L4*^ caused total loss of the proboscis suggesting for the first time an important interaction of *esg* and *dpp* that is specific for the labial disc, yet this interaction must happen in stages earlier that third instar, as we found that there is no overlap in the expression domains of *esg* and *dpp* in third instar labial discs.

The most affected structures by *esg* reduction were the antennae, maxillary palp, and proboscis which are the main *Drosophila* chemosensory organs. It is intriguing that the eye is barely affected in the *esg*^*L4*^ mutants, and this probably reflects the fact that the eye is more primitive and has different differentiation and developmental pathways than the antennae and the proboscis that are derived from primordial appendages. The different ontogenetic origins of these two structures is supported by the work of Gibson and Schubiger where they show that the overexpression of *fringe* (*fng*) using the same peripodial driver (C311-GAL4) that we used, only affects the eye development with no effect on the antennal and maxillary palps. We have previously shown that *esg*^*L4*^ was affected in chemosensory perception, the previously reported data and the phenotypes found in this work, suggest that *esg* function is critical for the determination of a morphogenetic field that induces the determination and development of the head chemosensory organs.

## Materials and methods

### Stocks and crosses

Flies were raised on standard yeast medium at 25°C or 28°C. *Oregon-R* and *w*^*1118*^ [Bloomington *Drosophila* Stock Center (BDSC)]. *L4* (*w*^*1118*^; *P{GawB-GAL4. esg}*^*L4*^) was obtained in our laboratory by mobilization of P-element P{GawB}. *UAS-iRNA esg* (Bloomington Stock # 34063), all other lines used in this work are available at the Bloomington Drosophila Stock Center (NIH P40OD018537). RNAi knockdown: *UAS-esg-RNAi* (BDSC 28514 and BDSC 34063), *UAS-esg-RNAi* [Vienna *Drosophila* Resource Center (VDRC), v9794 and v9793], *UAS-dpp-RNAi* (BDSC 25782). *C311-GAL4* (BDSC 5937), *dpp-lacZ* (BDSC 12379). *UAS-G-TRACE* (BDSC 28280). *P{EP}esg* (BDSC 30934)

### Immunohistochemistry

Tissue was dissected in PBS and fixed in PBS 4% formaldehyde and permeabilized with 0.2% TritonX-100 and blocked for 30 min with 5% goat pre-immune serum at room temperature. The primary antibodies were added in permeabilization buffer (see antibodies and concentrations below), incubated overnight at 4°C and washed 3 times in PBS 0.2% TritonX-100 at room temperature. The secondary antibodies were added, incubated for 90 min, and washed 3 times in PBS 0.2% TritonX-100 at room temperature. Primary antibodies used: Rabbit anti-GFP (Abcam ab290 1:1000), Rat anti-Elav [7E8A10, Developmental System Hybridoma Bank (DSHB) 1:50], Rat anti-*D*E-cadherin (DCAD2, DSHB 1:50), Mouse anti-Arm (N2 7A1, DSHB 1:50), Mouse Anti-En (4D9, DSHB 1:50), Rat anti-Ci (2A1, DSHB 1:50). Secondary antibodies Cy2 anti-rabbit, Cy2 anti-mouse, Cy2 anti-Rat, Cy3 anti-mouse, Cy3 anti-rat and Cy5 anti-mouse (Jackson immunoResearch) at 1:300 dilutions. Samples were mounted in Citifluor (Ted Pella Inc).

### Confocal Imaging

Images were collected at the “Laboratorio Nacional de Microscopía Avanzada (LNMA)” in an inverted Confocal multiphotonic Olympus FV1000 using an optimal confocal pinhole and Z steps. Images were analyzed and reconstructed using the open source ImageJ program (Schneider, Rasband and Eliceiri, 2012).

### TUNEL cell death detection

Cell death was labeled using the In Situ Cell Death Detection Kit, TMR red (Roche). Imaginal discs were dissected in PBS, fixed in PBS 4% paraformaldehyde for 1hr at 4°C, washed three time in PBST (PBS supplemented with 0.1% Triton X-100). Discs were blocked in PBST with 5% pre-immune goat serum for 12 hours at 4°C and transferred to permeabilization solution (0.1% Sodium citrate, 0.1% Triton X-100) at room temperature then followed the manufacturer’s instructions. Labeled were imagined using confocal microscopy. Signal was quantified using imageJ’s analyze particle tool.

### Quantitative PCR

qPCR reactions were performed using the universal TaqMan^®^ PCR master mix (Applied Biosystems) according to manufacturer instructions. However, the reactions were scaled down to a volume of 14 μl. The TaqMan^®^ probe was used at a final concentration of 25 nM, and 1 μl of a 1:20 dilution of the cDNA reaction was used for the detection 50 ng total, for each reaction of qPCR we used 5 ng. The TaqMan probe for *esg* used 5’-/56FAM/TTCTTGATGTCCGAGTGGGTCTGC/36-TAMSp/-31 that flanked by the primers 5’-TGCAGATCGCTCGAATCTGC-3’ and 5’-AGCAACTGGTGCAGGAGTAC-3’. The following PCR conditions were used: 95 °C for 10 min, 40 cycles of 95 °C for 15 s, and 58 °C for 1 min.

### Western blots

Third instar larvae were homogenized in homogenization buffer (sucrose 250 mM, Tris 50mM pH 7.4, KCl 25mM, MgCl_2_ 5mM, EDTA 5mM, 1mM Phenylthiourea, 2mM CaCl_2_, PMSF 1mM). Proteins were quantified using standard Bradford assay. Electrophoresis was performed in 7.5% polyacrylamide-SDS gels with 100 µg of total protein at 80 constant V. Proteins were transferred to 0.2 µm nitrocellulose membranes (# 1620112 BIO RAD) for 70 min at 120 V. Membranes were blocked in PBS, Tween 20 0.1%, milk 10% (Carnation Nestle) overnight at 4°C. Primary antibody was added (Anti-*D*E-cadherin (DCAD2, DSHB 1:1000), anti-Actin (JLA20, DSHB 1:1000)) in PBS, Tween 20 0.1%, milk 5% for 2 hours at room temperature. After three 15 minutes washes with PBST (PBS, Tween 20 0.1%) the secondary antibody (HRP-Anti Rat (Life Technologies) 1:4000 was added and incubated for two hours at room temperature. After three 15 minutes washes with PBST at room temperature signal was detected using the SuperSignal™ West Femto kit (Thermo Scientific, # 34095). Images were processed and quantified using ImageJ program.

## Acknowledgements

We thank René Hernández Vargas and Agustín Reyes for technical support, Santiago Becerra for oligonucleotide synthesis and Jorge Yañez for DNA sequencing, Ing. Roberto Pablo Rodríguez and David Santiago Castañeda of the “Unidad de cómputo of the Instituto de Biotecnología” for computer maintenance and technical support. We also thank Andrés Saralegui Amaro and Jaime Arturo Pimentel Cabrera from the Laboratorio Nacional de Microscopía Avanzada (LNMA) for support in confocal imaging.

**The authors declare no competing interests**

---

**Fig. S1.**
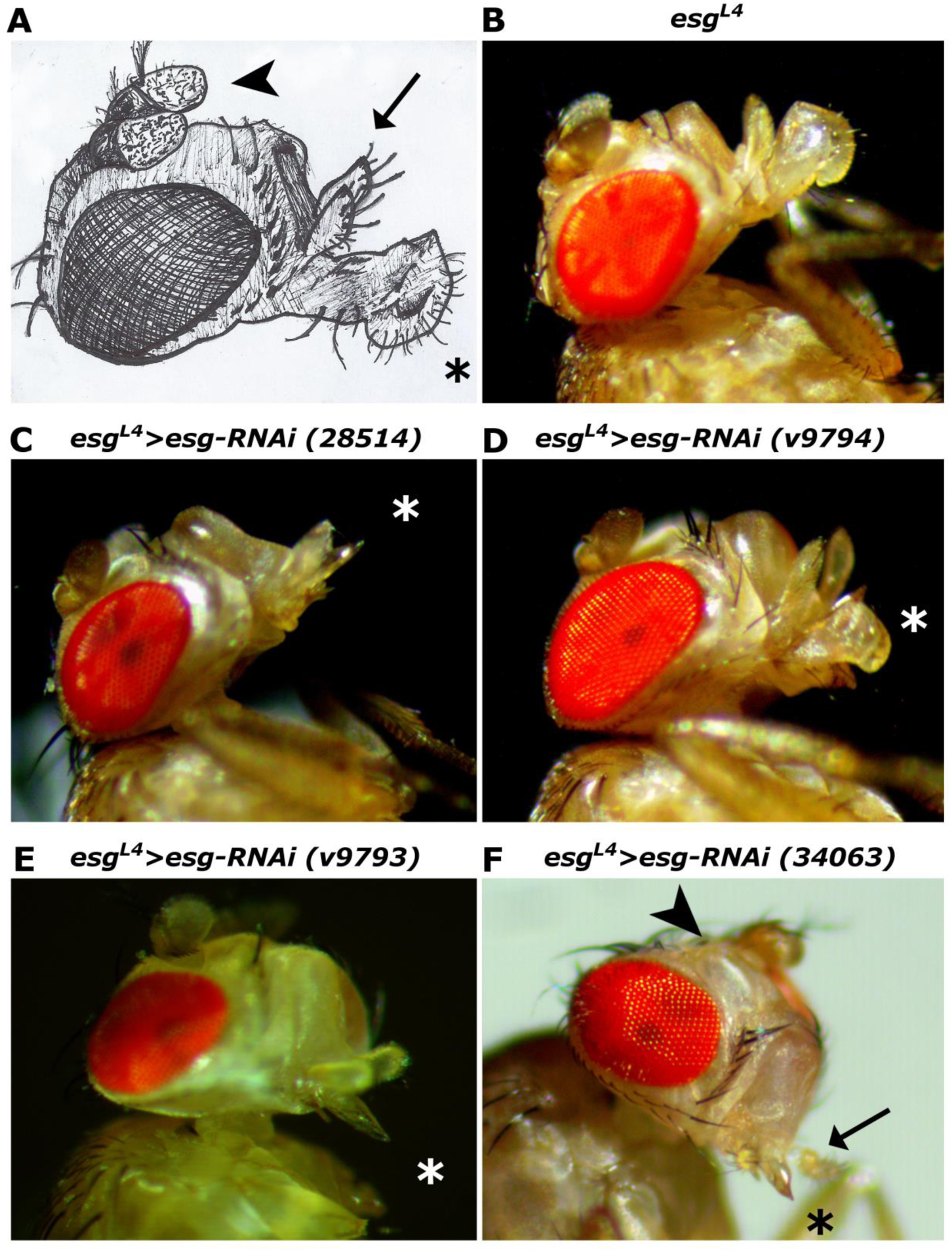
Phenotypes of the developmental defects in head structures induced by different RNAis targeting *esg*. **A)** Schematic representation of wt *Drosophila* head structures affected by the RNAis. **B-F)** Allelic series showing proboscis damage caused by the different anti *esg* RNAis driven by *esg*^L4^. **B)** *esg*^L4^ (control) shows fully developed proboscis and antennae. **C)** *esg*^*L4*^*>esg-RNAi(28514)* presents partial proboscis damage but no antennal defects. **D)** *esg*^*L4*^*>esg-RNAi(v9794)* all individuals present partial proboscis and antennal damage; in this example a single labellum is affected. **E)** *esg*^*L4*^*>esg-RNAi(v9793)* all individuals present complete loss of the proboscis from its base, only the labrum persists**. F)** *esg*^*L4*^*>esg-RNAi(34063)* has a lethal phenotype, the few individuals that reached the pupal stages were dissected to be analyzed; this genotype presents loss of antennae and complete loss of the proboscis from its base, only the labrum persists. Asterisks mark proboscis damage. Arrowhead points to defective maxillary palps. Arrow points to antennal damage.

**Fig. S2.**
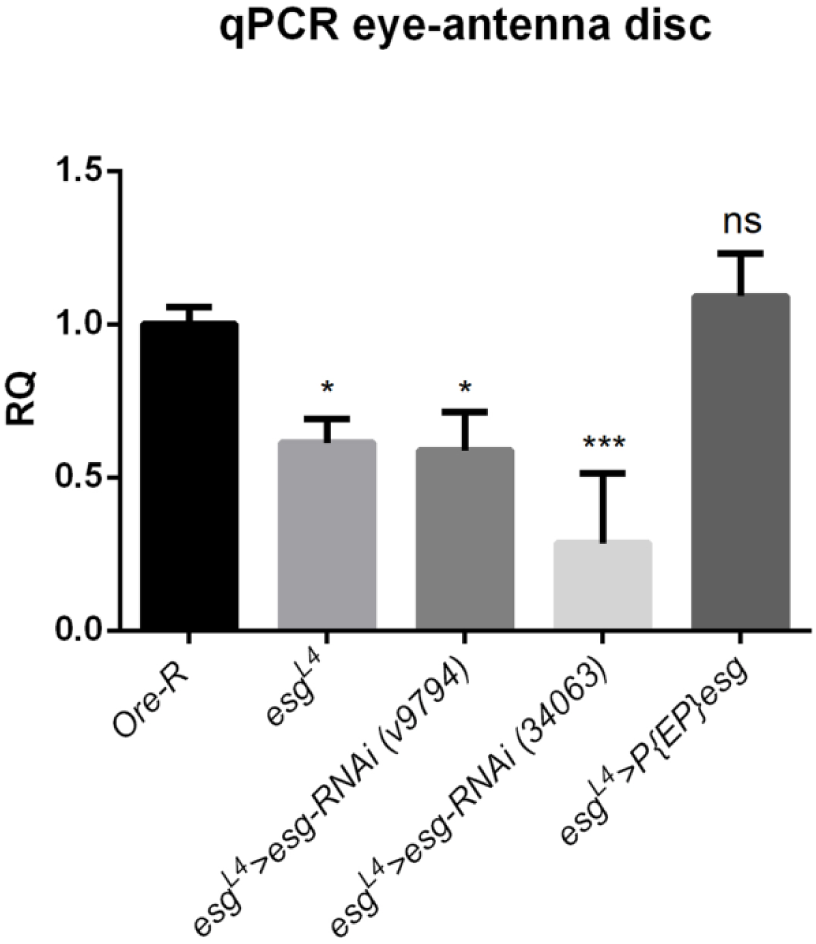
Quantitative PCR of *esg* levels from eye-antenna discs of different genotypes.

